# Shedding light on the Ophel biome: The Trans-Tethyan phylogeography of the sulfide shrimp *Tethysbaena* (Peracarida: Thermosbaenacea) in the Levant

**DOI:** 10.1101/2023.07.19.549641

**Authors:** Tamar Guy-Haim, Oren Kolodny, Amos Frumkin, Yair Achituv, Ximena Velasquez, Arseniy R. Morov

## Abstract

*Tethysbaena* are small peracarid crustaceans found in extreme environments such as subterranean lakes and thermal springs, represented by endemic species found around the ancient Tethys, including the Mediterranean, Arabian Sea, Mid-East Atlantic, and the Caribbean Sea. Two *Tethysbaena* species are known from the Levant: *T. relicta*, inhabiting the Dead Sea-Jordan Rift Valley, and *T. ophelicola*, found in the Ayyalon cave complex in the Israeli coastal plain, both belonging to the same species-group based on morphological cladistics. Along the biospeleological research of the Levantine subterranean fauna, three biogeographic hypotheses determining their origins were proposed: (1) Pliocenic transgression, (2) Mid-late Miocenic transgression, and (3) The Ophel Paradigm, according to which these are inhabitants of a chemosynthetic biome as old as the Cambrian. We have used mtDNA COI gene and a molecular clock approach to establish the phylogeny and assess the divergence times of the Levantine *Tethysbaena*. Contrary to prior hypotheses, our results indicate a two-stage colonization pattern: a late Oligocene transgression, through a marine gulf extending from the Arabian Sea, leading to the colonization of *T. relicta* in the Dead Sea-Jordan Rift Valley, and a Miocene transgression in the emerging Mediterranean region, carrying *T. ophelicola* to the coastal plain of Israel.

## INTRODUCTION

Groundwater fauna (stygofauna) is characterized by short-range endemism and high species crypticity. The unique suite of troglomorphic traits (e.g., loss of pigment, reduced eyes) characterizing stygobionts often hinders distributional studies due to the highly convergent characteristics that can obscure taxonomic relationships (Juan, Guzik, Jaume & Cooper, 2010; Porter, 2007). As a result, molecular phylogenetic tools have been extensively used over the last two decades to infer stygofauna biogeographies and the underlying processes shaping them (e.g., Abrams, Huey, Hillyer, Humphreys, Didham & Harvey, 2019; Asmyhr, Hose, Graham & Stow, 2014; Bauzà-Ribot, Juan, Nardi, Oromí, Pons & Jaume, 2012; Bradford, Adams, Humphreys, Austin & Cooper, 2010; Cánovas, Jurado[Rivera, Cerro[Gálvez, Juan, Jaume & Pons, 2016; Cooper, Fišer, Zakšek, Delić, Borko, Faille & Humphreys, 2023; Finston, Bradbury, Johnson & Knott, 2004; Guy-Haim, Simon-Blecher, Frumkin, Naaman & Achituv, 2018; Jaume, 2008; Jurado-Rivera, Pons, Alvarez, Botello, Humphreys, Page, Iliffe, Willassen, Meland & Juan, 2017; Marin, Krylenko & Palatov, 2021; Matthews, Abrams, Cooper, Huey, Hillyer, Humphreys, Austin & Guzik, 2020).

Thermosbaenacea is a small order of peracarid crustaceans comprising unique and highly specialized species adapted to extreme aquatic environments, including spring-fed subterranean lakes and thermal springs, with their core populations found deep underground in the inaccessible phreatic waters. Anoxic, sulfide-rich environments are favorable to Thermosbaenacea—often feeding on bacterial mats formed by sulfide-oxidizing bacteria— thus termed “sulfide shrimp” by Por (2014). Based on their distribution, it was assumed that the ancestral habitat of the thermosbaenaceans is the ancient Tethys Sea, and they are represented by relic fauna found around the Mediterranean, the Arabian Sea, Mid-East Atlantic, and the Caribbean Sea (Hou & Li, 2018; Wagner, 1994). Among thermosbaenaceans, *Tethysbaena* (family: Monodellidae) is the most speciose and widespread genus, comprising 27 species in seven species-groups (Wagner, 1994; Wagner & Bou, 2021). Only a few of the *Tethysbaena* species-groups were analyzed and supported by molecular phylogenetic tools (Cánovas et al., 2016; Wagner & Chevaldonné, 2020).

Two species of *Tethysbaena* are known from Israel: *T. relicta* Por, 1962 (formerly *Monodella relicta*) and *T. ophelicola* Wagner, 2012. Initially, fragments of *T. relicta* were found in the hot spring Hamei Zohar by the Dead Sea in Israel (Por, 1962). Later, scattered specimens of the same species were collected from the thermohaline spring En-Nur, on Lake Kinneret shore, a few hundred kilometers to the north (Dimentman & Por, 1991), thus inferring that *T. relicta* inhabits the whole groundwater system of the Dead Sea-Jordan Rift Valley aquifer. *T. ophelicola* was found in the karstic underground basin near Ramla, named Ayyalon-Nesher-Ramla complex (Por, 2014; Por, Dimentman, Frumkin & Naaman, 2013; Wagner, 2012), 60 km west of the Dead Sea-Jordan Rift Valley, beyond the water divide of Israel.

Based on synapomorphies of the antennular inner flagellum and maxilliped macrosetae (Wagner, 1994), it was hypothesized that together with other closely allied species—one species from Somalia (Chelazzi & Messana, 1982), four species from Oman (Wagner, 2020), one species from Yemen (Wagner & Van Damme, 2021)—*T. relicta* and *T. ophelicola* form the “*T. relicta*-group” (Wagner, 2012), suggesting a recent common ancestor. An alternate hypothesis can be drawn from the phylogenetic analysis of the prawn *Typhlocaris* (Guy-Haim et al., 2018), preying on *Tethysbaena* in Ayyalon and En-Nur (Tsurnamal, 1978; Tsurnamal, 2008; Tsurnamal & Por, 1971; Wagner, 2012). Four *Typhlocaris* species are known, two of which co-occur with *Tethysbaena*: *Ty. galilea* inhabiting En-Nur spring (Calman, 1909; Tsurnamal, 1978) and *Ty. ayyaloni* from the Ayyalon cave (Tsurnamal, 2008). The two additional *Typhlocaris* species are *Ty. salentina* from Apulia region in Southeastern Italy (Caroli, 1923; Froglia & Ungaro, 2001) and *Ty. lethaea* from Libya near Benghazi (Parisi, 1921). The molecular phylogeny of *Typhlocaris* species showed that *Ty. ayyaloni* (Israel) and *Ty. salentina* (Italy) are more closely related to each other than either of them is to *Ty. galilea* (Israel) (Guy-Haim et al., 2018). Accordingly, we can hypothesize a similar phylogeographic pattern of the Levantine *Tethysbaena*, where *T. ophelicola* would be more closely related to the Mediterranean species (“*T. argentarii*-group”) than to *T. relicta*.

Along the biospeleological research of the Thermosbaenacea and other phyla of subterranean crustaceans represented in the Dead Sea Rift Valley (Syncarida, and the families Bogidiellidae and Typhlocarididae), three paradigms have been proposed to explain their origins: (1) Pliocenic marine transgression (Por, 1963), (2) Miocenic Tethys transgression (Dimentman & Por, 1991; Por, 1987), and (3) The Ophel Paradigm that offered a conceptual framework, within which these styobionts are inhabitants of the ancient chemosynthetic Ophel biome, dating back at least to the Cambrian (Por, 2011). Using a molecular clock approach, Guy-Haim et al. (2018) estimated the divergence time of the *Typhlocaris* species. They based their analysis on a calibration node inferred from a regional geological event— the end of the marine connection between the Mediterranean Sea and the Dead Sea-Jordan Rift Valley, marked by the top of Bira formation, dated to 7 MYA (Rozenbaum, Sandler, Zilberman, Stein, Jicha & Singer, 2016), separating *Ty. galilea* and the *Typhlocaris* ancestor. The inferred divergence time of *Ty. ayyaloni* and *Ty. salentina* was 5.7 (4.4–6.9) MYA, at the time of the Messinian Salinity Crisis (5.96–5.33 MYA), when the Mediterranean Sea desiccated and lost almost all its Miocene tropical fauna (Por, 1987; Por, 1989). It is therefore an open question as to whether the same vicariant events have shaped the biogeographies of both predator (*Typhlocaris*) and prey (*Tethysbaena*) subterranean crustaceans.

The main objectives of our study were to (1) reveal the phylogenetic relatedness of the Levantine *Tethysbaena* species, and use these patterns to (2) infer the geological and evolutionary processes that have shaped their divergence patterns.

## MATERIALS AND METHODS

### Sampling sites, specimen collection and identification

Specimens of *T. ophelicola* were collected by a hand pump from the inner pool of the Levana cave (31.9223°N, 34.8942°E), part of the Ayyalon-Nesher-Ramla complex (Fig. 1).

**Figure 1.**
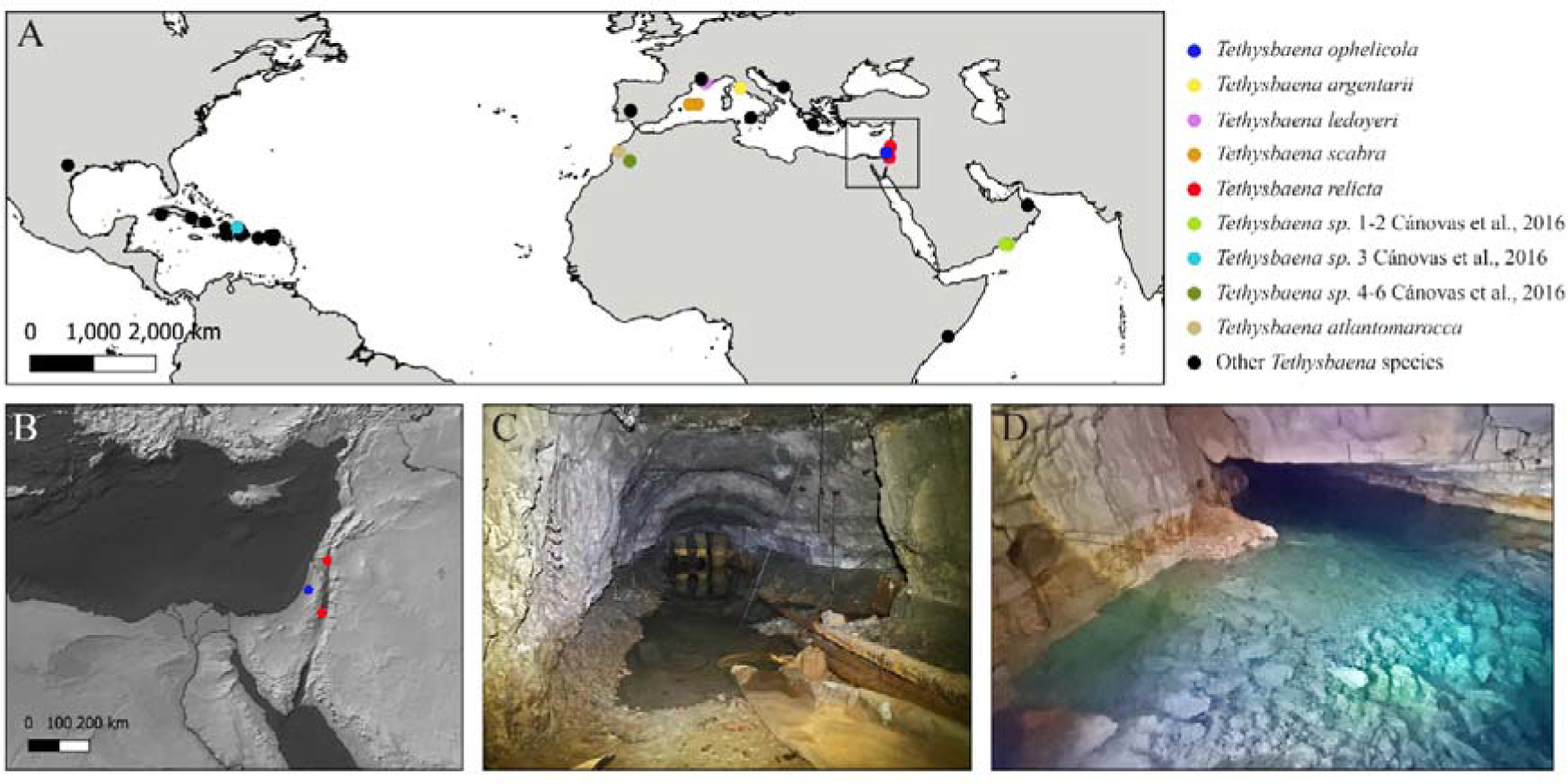
*Tethysbaena* distribution and habitats. **A**. Global *Tethysbaena* distribution. The species included in the phylogenetic analysis are presented in colored circles. Other *Tethysbaena* species are presented in black. Based on documented records in Wagner (1994); Wagner (2012); Wagner (2020), Cánovas et al. (2016); Wagner and Chevaldonné (2020) and Wagner and Bou (2021). **B.** Levantine distribution of *T. ophelicola* and *T. relicta*. **C-D**. *Tethysbaena* Levantine habitats. **C**. An artificial tunnel near the Dead Sea, Israel. **D**. Levana (Ayyalon) cave, Israel.

Specimens of *T. relicta* were collected by a hand pump from an artificial tunnel near the Dead Sea Shore penetrating the Judea Group aquifer, 6.5 km north of Hamei-Zohar (31.2232°N, 35.3547°E) (Fig. 1). The *locus typicus* of *T. relicta*, the thermal spring of Hamei-Zohar (Por, 1962), is no longer accessible since the 1970s, as hotels were built on the spring area.

Part of the collected specimens was preserved in 70% ethanol and the other in absolute ethanol for morphological and molecular analyses, respectively. Species identification of *T. ophelicola* and *T. relicta* was performed using a stereomicroscope (SZX16, Olympus, Japan) following the identification keys in Por (1962) and Wagner (1994); Wagner (2012).

### DNA extraction, amplification and sequencing

Cánovas et al. (2016) used both mitochondrial cytochrome *c* oxidase subunit I (COI) and nuclear 28S rRNA genes to assess the genetic population structure of the anchialine *T. scabra* in the Balearic Islands, and found that the 28S rDNA gene showed low genetic variation resulting in a poorly resolved phylogenetic tree, and they, therefore, based their phylogenetic reconstruction and divergence time estimations on the COI gene only. Following their finding, we have used the COI gene in our analysis.

Total genomic DNA was extracted from each individual using the DNEasy Blood and Tissue kit (QIAGEN, Germany) according to the manufacturer’s specifications. Following the DNA extraction, the COI gene was amplified using PCR with universal primers LCO1490 and HCO2198 (Folmer, Black, Hoeh, Lutz & Vrijenhoek, 1994). Reaction conditions were as follows: 94 °C for 2 min, followed by 5 cycles of 94 °C for 40 s, 45 °C for 40 s, and 72 °C for 1 min, and followed by 30 cycles of 94 °C for 40 s, 51°C for 40 s, and 72 °C for 1 min, and a final elongation step of 72 °C for 10 min. Obtained PCR products were purified and sequenced by Hylabs (Rehovot, Israel).

### Phylogenetic analysis

A total of 22 COI sequences of *Tethysbaena* were analyzed, including *T. ophelicola* (n=3) and *T. relicta* (n=3) obtained in this study. Additional sequences of *T. scabra* (Balearic Islands, n=5), *T. argentarii* (Italy, n=2), *T. ledoyeri* (France, n=2), *T. atlantomaroccana* (Morocco, n=1), and further sequences of *Tethysbaena* sp., unidentified to the species level, from Oman (n=2), Morocco (n=3) and the Dominican Republic (n=1), were obtained from NCBI GenBank (https://www.ncbi.nlm.nih.gov/genbank/) and the European Nucleotide Archive (https://www.ebi.ac.uk/ena/browser/home). The thermosbaenacean *Halosbaena tulki* was chosen as an outgroup following Page, Hughes, Real, Stevens, King and Humphreys (2016) and used as a root node in the phylogenetic analysis. All specimens, collection sites, accession numbers, and related references are summarized in Table 1.

**Table 1:**
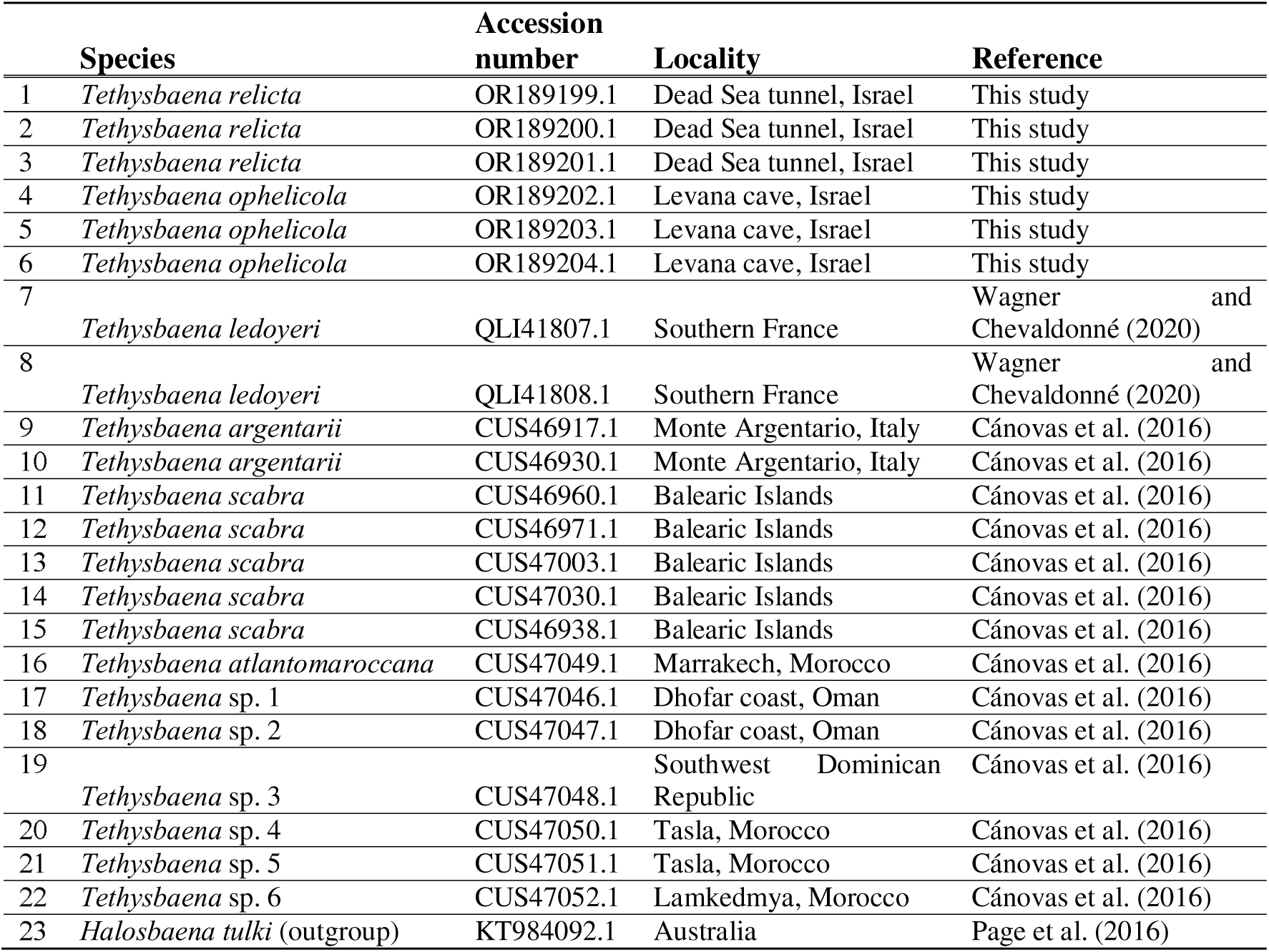
*Tethysbaena* species and outgroup included in the phylogenetic analysis.

Sequence alignment was conducted using ClustalW embedded in MEGA v11.0 (Tamura, Stecher & Kumar, 2021). The best-fitting substitution model was selected according to the Bayesian Information Criterion using Maximum-likelihood (ML) model selection in MEGA. GBlocks v0.91.1 (Castresana, 2000) was used for trimming the ambiguous blocks in the sequence alignment. ML analysis was performed using the T92+G+I model (BIC= 6112.5) with 1000 bootstrapping replicates. Bayesian Metropolis coupled Markov chain Monte Carlo (B-MCMC) analyses were conducted with MrBayes v3.2.7a (Ronquist, Teslenko, Van Der Mark, Ayres, Darling, Höhna, Larget, Liu, Suchard & Huelsenbeck, 2012) on XSEDE in the CIPRES v3.3 Science Gateway portal (https://www.phylo.org/portal2) with nst=2, rates=gamma, and statefreqpr=fixed(fixedest=equal). Two independent runs of 10,000,000 generations each performed, sampling every 1000 generations. A burn-in at 25% of the sampled trees was set for final tree production. Convergence and effective sampling of runs was assessed using Tracer v. 1.6 (Drummond & Rambaut, 2007), and the post-burnin tree samples were summarized using the sumt.

### Estimation of divergence times

Molecular clock calculations for cave-dwelling species are often contentious (Page, Humphreys & Hughes, 2008). Stygobionts often exhibit unique evolutionary characteristics and experiences, including isolation, reduced gene flow, small population sizes, and distinct selective pressures. These factors can lead to deviations from a constant rate of molecular evolution among lineages, rendering a strict molecular clock assumption less realistic. Therefore, we used a relaxed molecular clock approach (Drummond, Ho, Phillips & Rambaut, 2006). Cánovas et al. (2016) assessed the divergence time of the Western Mediterranean *Tethysbaena*, *T. scabra* from the Balearic Islands, and *T. argentarii* from Italy using the COI gene. They based the substitution rates on the mean rate estimated for a co-occurring anchialine stygobiont amphipod *Metacrangonyx longipes*, 1.32% per lineage and million years (0.89–1.95, 95% CI) (Bauzà-Ribot et al., 2012). Following Cánovas et al. (2016), we implemented this substitution rate in our dataset.

A relaxed-clock MCMC (Markov Chain Monte Carlo) approach using the uncorrelated log-normal model was implemented in BEAST v2.4 (Drummond & Rambaut, 2007; Suchard, Lemey, Baele, Ayres, Drummond & Rambaut, 2018; Suchard & Rambaut, 2009). The Yule process was chosen as speciation process. Three independent runs, each of 50,000,000 generations, were performed, with sampling every 5000 generations. The three separate runs were then combined (following the removal of 10% burn-in) using LogCombiner v2.5.2. Log files were analyzed with Tracer v1.6 (Drummond & Rambaut, 2007), to assess convergence, confirm the combined effective sample sizes for all parameters, and ensure that the MCMC chain had run long enough to get a valid estimate of the parameters (Drummond & Rambaut, 2007). Maximum clade credibility (MCC, hereafter) tree was then produced using TreeAnnotator v2.4.7 (Rambaut & Drummond, 2017). FigTree v.1.4.4 (Rambaut, 2018) was used to visualize the MCC tree and the highest posterior density (HPD, hereafter) ranges.

## RESULTS

### Morphological identification

Specimens of *T. relicta* collected from the Dead Sea tunnel were similar to the specimens from Hamei-Zohar thermal spring described by Por (1962), and included males, with no ovigerous or brooding females (Fig. 2A). The average length (excluding antennae) was 2104±181 µm (n=5, ±SD, hereafter). The following morphological features characterized the specimens as belonging to *T. relicta*: 8 segments in the main flagellum (endopodite) of antenna 1; 7 terminal plumidenticulate macrosetae on the maxilliped; the uropod included 5 medial plumose macrosetae, 11–13 plumose macrosetae in the endopodite, and 16–19 macrosetae in the second segment of the exopodite. The mean width:length ratio of the telson was 1.15.

**Figure 2.**
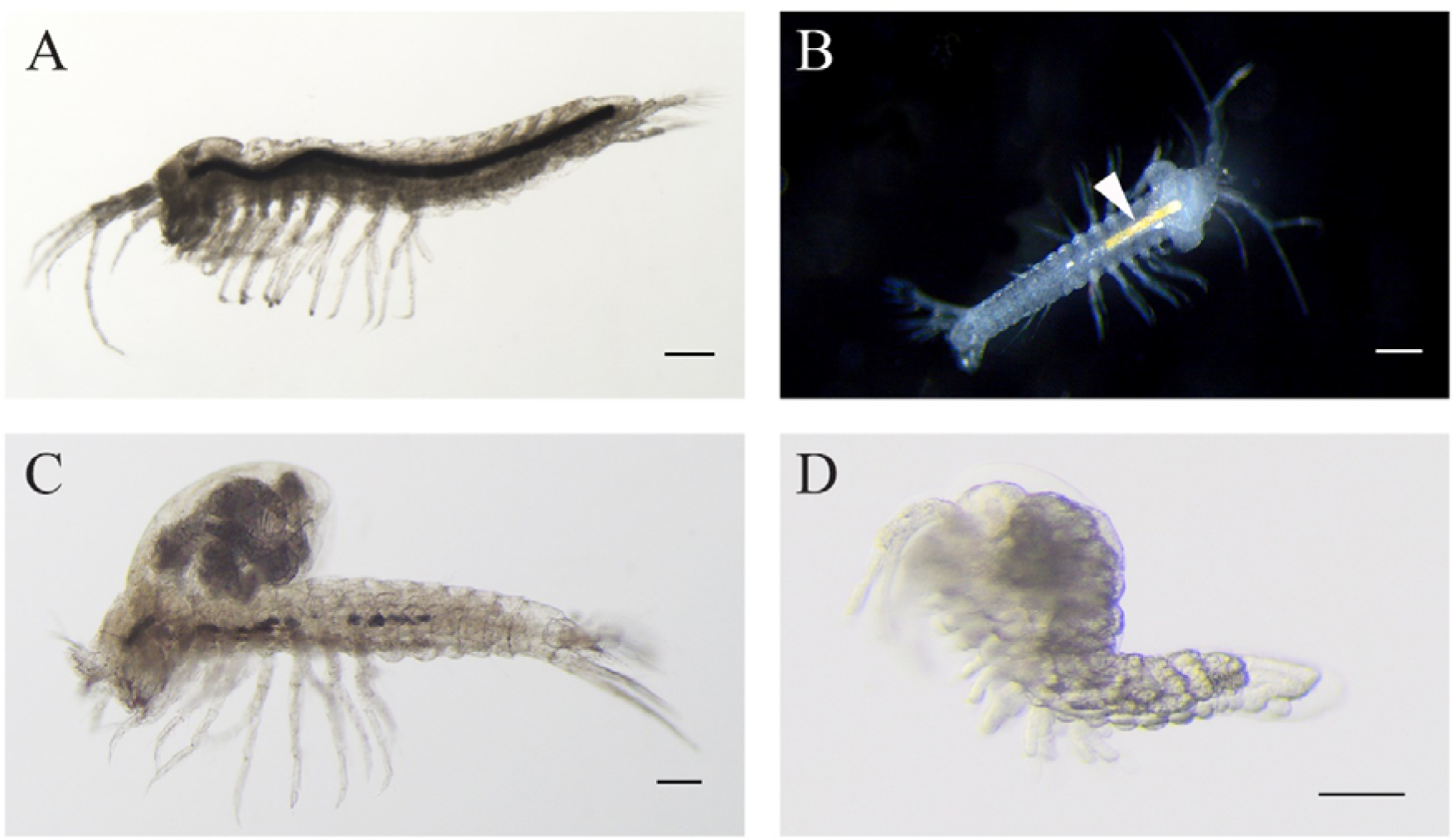
**A**. *Tethysbaena relicta* Por, 1962 male. **B-D**. *Tethysbaena ophelicola* Wagner, 2012 male (B), brooding female (C), and postmarsupial juvenile (D). The arrowhead points to the orange coloration of the gut (B), indicating the presence of sulfide-oxidizing bacteria. The scale bar denotes 200 µm in A-C and 100 µm in D.

Specimens of *T. ophelicola* from Levana cave were similar to the specimens from Ayyalon cave described by Wagner (2012), and included males, brooding females and postmarsupial juveniles (Fig. 2B–D). The average length (excluding antennae) was 2276±380 µm in males (n=5) and 2620±139 µm in females (n=5). The following morphological features were found: 7 segments in the main flagellum (endopodite) of antenna 1; 7 terminal plumidenticulate macrosetae on the maxilliped; uropod included 4 medial plumose macrosetae and 18–22 plumose macrosetae in the endopodite and the second segment of the exopodite. The mean width:length ratio of the telson was 1.10.

### Molecular phylogenetic analysis

The DNA barcode consisting of a fragment of 708 bp of the COI gene was sequenced from 6 specimens of *T. ayyaloni* and *T. relicta*. Sequences were deposited in NCBI GenBank under accession numbers OR189199–OR189204. The phylogenetic analysis included 16 additional *Tethysbaena* sequences and one *Halosbaena tulki* sequence as an outgroup (Table 1). The overall alignment was 691 bp long, with 227 parsimonious informative sites.

ML and Bayesian phylogenetic analyses showed similar tree topologies (Fig. 3). The Levantine *Tethysbaena* species from Israel present polyphyly, where *T. ayyaloni* lies within a Mediterranean clade (including *T. scabra* from the Balearic Islands, *T. ledoyeri* from Southern France and *T. argentarii* from Italy) with 100% bootstrapping support and 0.99 posterior probability, and *T. relicta* clusters with *Tethysbaena* sp. from Oman (100% bootstrapping support and 1.00 posterior probability), and the Dominican Republic (87%/0.83 bootstrapping support/posterior probability), forming the Arabian-Caribbean clade. The Atlantic *Tethysbaena T. atlantomaroccana* is a sister taxon to the Mediterranean clade species, although with a lower support/probability. The other Moroccan *Tethysbaena* species from Tasla and Lamkedmya were in a more basal position but showed lower bootstrapping support (<50%).

**Figure 3.**
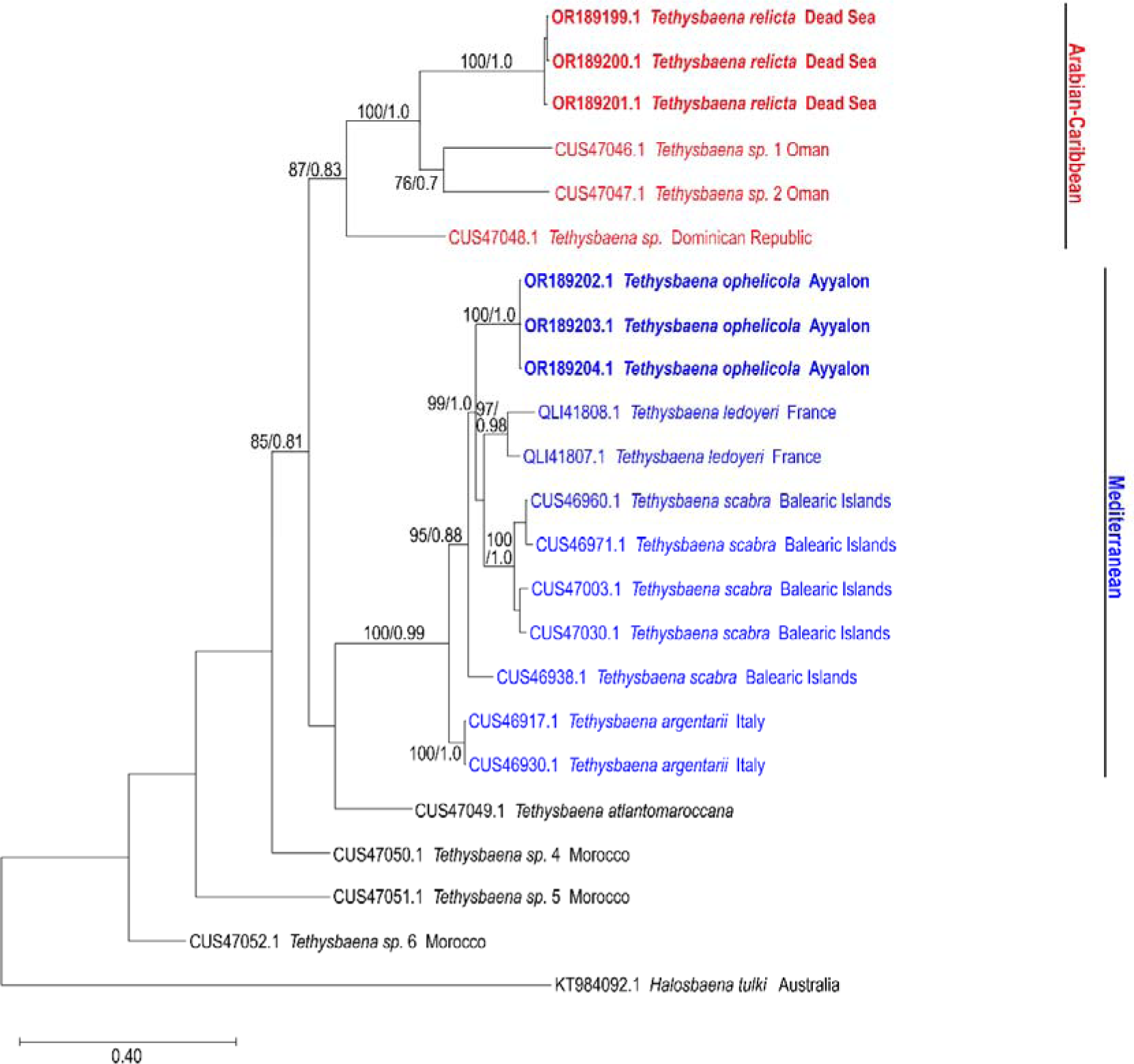
Maximum-Likelihood phylogenetic tree of *Tethysbaena* based on the COI gene, using the T92+G+I substitution model. *Halosbaena tulki* was used as a root node. At each node, the number on the left size of the slash indicates the percentage of ML bootstrap support (1000 replicates), and the right number indicates the Bayesian posterior probability expressed as a decimal fraction, for nodes that received at least 50% support. The scale bar denotes the estimated number of nucleotide substitutions per site.

**Figure 4.**
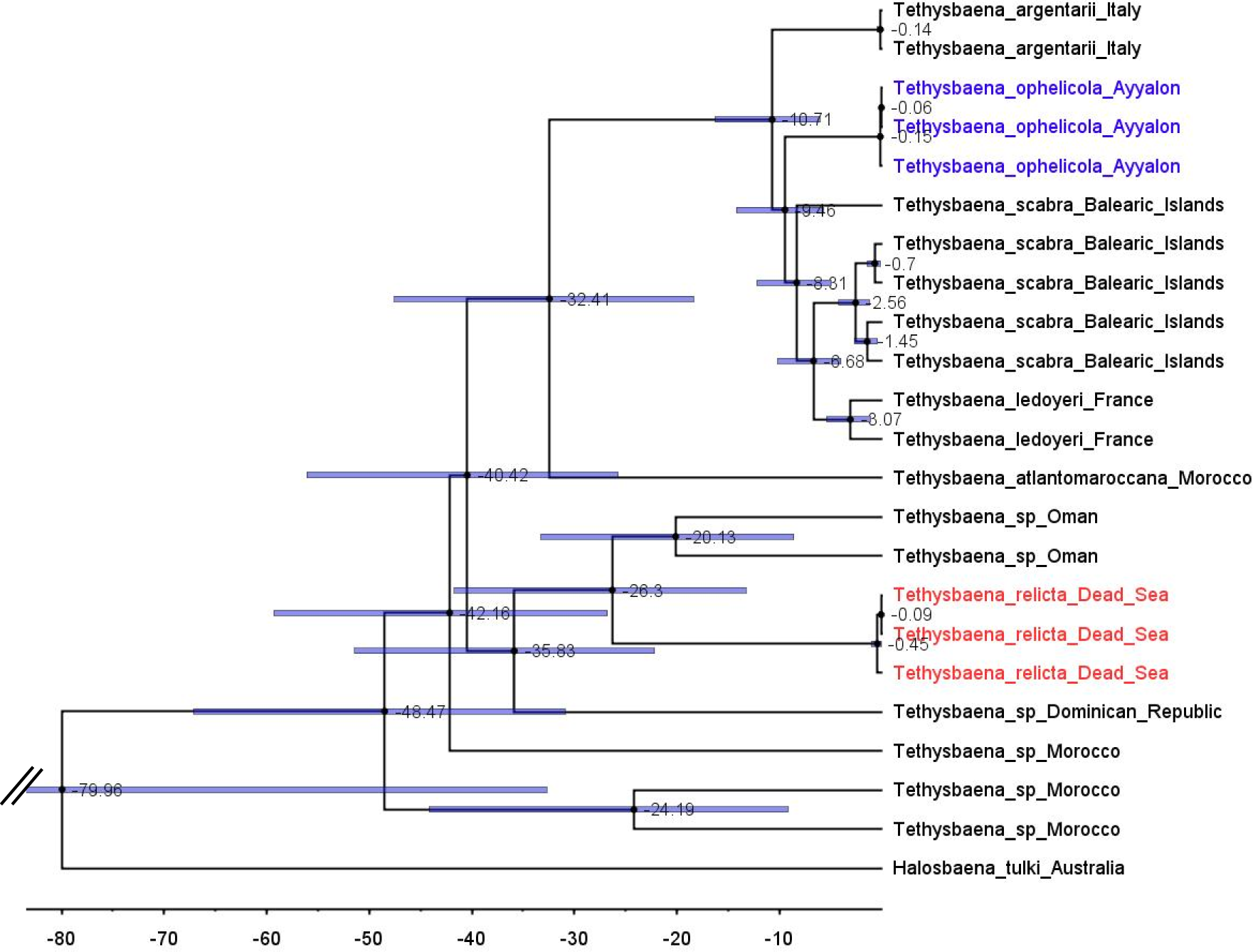
*Tethysbaena* time tree using the COI gene. A relaxed MCMC clock using the uncorrelated log-normal model and substitution rate based on Cánovas et al. (2016) were implemented in BEAST v2.4. Mean ages are presented on the nodes, and the 95% HPD (highest posterior density) are presented by the blue bars.

### Divergence time estimation

Effective sample size (ESS) values were at least 436 and 356 for the posterior and prior statistics, respectively, 1738 for the likelihood statistic, and greater than 1400 for all MRCA times estimates, suggesting good mixing and an effective MCMC sampling of the posterior distribution.

We estimated the ages for eight nodes (Table 2). The youngest node was the most recent common ancestor of *T. leyoderi* from Southern France and *T. scabra* from the Balearic Islands, which returned a mean estimate at 8.31 MYA with 95 % HPD of 10.15–3.97 MYA. The next mean estimate is the divergence of *T. ophelicola* from the clade of *T. leyoderi* and *T. scabra*, dated to 9.46 MYA, with 95% HPD of 14.20–5.71 MYA. The mean age of the most common ancestor of all Mediterranean *Tethysbaena* was 10.71 MYA with 95 % HPD of 16.27–6.04 MYA. The most recent ancestor of the Mediterranean clade and *T. atlantomaroccana* from Morocco was dated to 32.41 MYA with 95 % HPD of 47.53–18.37 MYA. The mean age of the node linking *T. relicta* from the Dead Sea-Jordan Rift Valley and *Tethysbaena* from Oman was 20.13 MYA with 95 % HPD of 41.69–13.25 MYA. The node of the most recent common ancestor of *T. relicta*, *Tethysbaena* from Oman, and the *Tethysbaena* from the Dominican Republic had a mean estimate of 35.84 MYA with 95 % HPD of 51.41–22.16 MYA. The mean age for the node linking the Arabian-Caribbean clade (*T. relicta* + *Tethysbaena* sp. Oman + *Tethysbaena* sp. Dominican Republic) with the Mediterranean-Atlantic clade (*T. scabra + T. leyoderi* + *T. ophelicola + T. argentarii* + *T. atlantomaroccana*) was 40.42 MYA with 95 % HPD of 56.09–25.72 MYA. The estimate for the root node linking *Tethysbaena* and *Halosbaena* was 79.96 MYA with 95% HPD of 137.8–32.68 MYA.

**Table 2:**
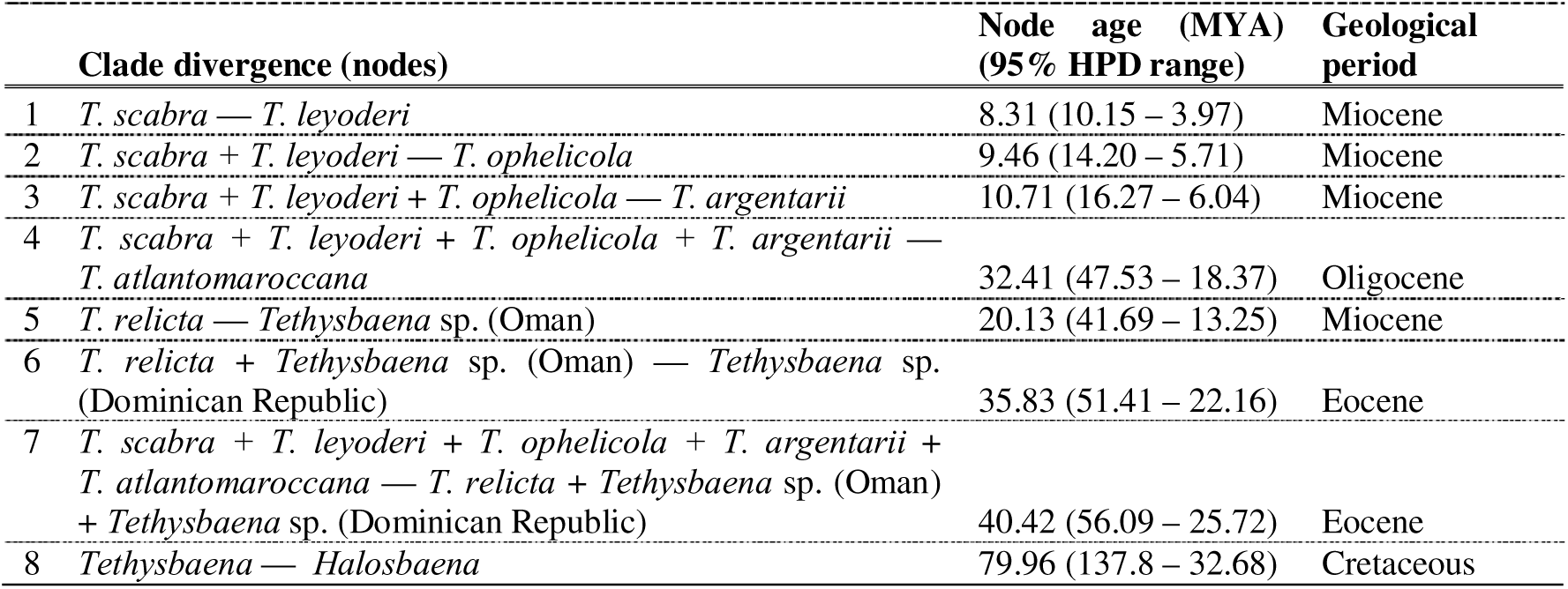
Divergence times for *Tethybaena* species as estimated by the Bayesian evolutionary analysis method calculated using the COI gene molecular evolution based on Cánovas et al. (2016) and Bauzà-Ribot et al. (2012). Node ages and highest posterior density (±95% HPD) ranges are given in million years round.

## DISCUSSION

In his monography on Thermosbaenacea, Wagner (1994) divided the Monodellidae family to two genera, the monotypic *Monodella* and the speciose *Tethysbaena*, which he named after the ancient Tethys Sea and the Greek word “βαινειν” (meaning “to walk”), referring to these animals as “walkers of the Tethys Sea”. He noted that although there is a great similarity among the different species, six species-groups can be identified based on morphological characters. With the later finding of *T. exigua* from Southern France, a seventh group was established (Wagner & Bou, 2021). Here, we analyzed the phylogenetic relatedness and divergence times of the two Levantine *Tethysbaena* species found in Israel: *T. relicta* from the Dead Sea-Jordan Rift Valley, and *T. ophelicola*, from the Ayyalon-Nesher-Ramla cave complex in central Israel.

According to Wagner (2012) and Wagner and Van Damme (2021), both Levantine species belong to “*T. relicta*-group” (together with four species from Oman, one species from Somalia and one species from Yemen), implying that these are sister taxa sharing a most recent common ancestor. Our results reject the morphology-based cladistics and support the hypothesis suggesting that *T. relicta* shared an ancestor with *Tethysbaena* species from Oman and Dominican Republic, whereas the circum-Mediterranean species (including *T. ophelicola*) share another ancestor. Indeed, discrepancies between morphological cladistics and molecular phylogeny are common in cave fauna and were often attributed to their troglomorphic traits (Bishop & Iliffe, 2012; Juan et al., 2010; Porter, 2007).

Three paradigms determining the origin of the Thermosbaenacea and other phyla of subterranean crustaceans represented in the Dead Sea-Jordan Rift Valley (Syncarida, and the families Bogidiellidae and Typhlocarididae) and around the Mediterranean were defined. The earlier paradigm suggested that the Levantine *Tethysbaena*, among other subterranean salt-water fauna, have resulted from a late Pliocenic pre-glacial (Piacenzian) marine transgression (Fryer, 1964; Hubault, 1937; Por, 1963). A narrow gulf penetrated into the coastal line near the present-day mount Carmel and then bent southwards along the Dead Sea-Jordan Rift Valley reaching a basin that extended south of the present Dead Sea (Picard, 1943). According to this paradigm, the Pliocenic Mediterranean was still inhabited by a very large number of Tethys remnants, including thermosbaenaceans, that were stranded in the Rift Valley and around the Mediterranean.

Por (1986) rejected the first paradigm, noting that the Pliocenic Mediterranean no longer contained the tropical fauna that include the *Tethysbaena* ancestor and that the short-lived Pliocenic transgression did not establish viable marine environments. Instead, he posited that these species represent marine fauna colonized by Miocenic transgression, the last time that tropical sea penetrated inland in the Levant, and left stranded following a late Miocene regression (Dimentman & Por, 1991; Por, 1987; Por, 1989). This second paradigm was supported by Guy-Haim et al. (2018) who used a molecular clock approach to estimate the divergence time of the *Typhlocaris* species, based on a calibration node inferred from the end of the marine connection between the Mediterranean Sea and the Dead Sea-Jordan Rift Valley, marked by the top of Bira formation, dated to 7 MYA (Rozenbaum et al., 2016). They inferred a divergence time of *Typholocaris* from Ayyalon cave and Italy of 5.7 (4.4– 6.9) MYA, at the time of the Messinian Salinity Crisis. During this event, the African plate moved towards the Euro-Asian plate, closing the Straits of Gibraltar and temporarily isolating the Mediterranean Sea from the Atlantic Ocean (Krijgsman, Hilgen, Raffi, Sierro & Wilson, 1999). As a result, the Mediterranean Sea partly desiccated and transformed into small hypersaline basins, losing almost all its Miocenic tropical fauna, including those able to colonize subterranean waters (Por, 1975; Por, 1986; Por, 1987; Por, 1989).

With the discovery of the Ayyalon cave system and its endemic stygofauna in 2006, a third paradigm known as “the Ophel Paradigm” was developed by Por (2007). He identified the “Ophel” as a continental subterranean biome, subsisting on chemoautotrophic bacterial food, independently of the exclusive allochthonous epigean food of photoautotrophic origin. Within this biome, *Tethysbaena* are primary consumers, presenting a typical feeding behavior of upside-down swimming-gathering of sulfur bacteria or bacterial mats (Por, 2011; Wagner, 2012). Following the development of the new chemosynthetic-based biome paradigm, Por presented an alternative to the Tethys stranding paradigm, stating that the “*Ophel paradigm falsified first of all my own, previously held views*” on the diversification of the subterranean fauna in the Levant (Por, 2011). He noted that the pre-Messinian fauna of the fossiliferous taxa of the foraminiferans, the mollusks and the teleost fishes was similar to the recent Red Sea fauna or different only at the species level, and there is no indication for extinction of crustaceans during the Tertiary, thus the origin of the subterranean Levantine fauna is of earlier origin (Por, 2010). Por suggested that the Ophelic biome is possibly at least as old as the Cambrian, which had a diverse aquatic crustacean and arthropodan palaeofauna, including Thermosbaenacea (Por, 2011).

Cánovas et al. (2016) assessed the divergence time of the Western Mediterranean *Tethysbaena*, *T. scabra* from the Balearic Islands and *T. argentarii* from Italy using the COI gene. They based the substitution rates on the mean rate estimated for a co-occurring anchialine stygobiont amphipod *Metacrangonyx longipes*, 1.32% per lineage and million years (0.89–1.95, 95% CI) (Bauzà-Ribot et al., 2012) and estimated the divergence time of *T. scabra* and *T. argentarii* to the early Tortonian, 10.7 MYA. Following Cánovas et al. (2016), we have used the COI gene to assess the divergence times of the Levantine *Tethysbaena*, *T. relicta* and *T. opehlicola*, and additional *Tethysbaena* species from around the Mediterranean, Arabian, and Caribbean Sea, using the Australian *Halosbaena* as an outgroup.

Our analysis shows that the divergence times of *Tethysbaena* species are earlier than those of *Typhlocaris* species, pre-dating the upper-Miocene Messinian Salinity Crisis. Most divergence events occurred in the Miocene and Oligocene. The Dead Sea-Jordan Rift Valley *T. relicta* shares a most recent common ancestor with *Tethysbaena* from the Arabian Sea (Oman), dated to the early Miocene, 20.13 MYA (with 95% HPD of 41.69 – 13.25), corresponding with the Oligo-Miocene rift-flank uplift of the Arabian plate during the formation of the Red Sea and Gulf of Aden (Omar & Steckler, 1995; Stern & Johnson, 2010). Both *T. relicta* and the *Tethysbaena* from Oman separated from the Caribbean *Tethysbaena* during the Eocene-Oligocene transition, when global cooling and tectonic uplift caused sea level decline and led to the establishment of the modern Caribbean Seaway (Iturralde-Vinent & MacPhee, 1999; Iturralde-Vinent, 2006; Weaver, Cruz, Johnson, Dupin & Weaver, 2016).

The most recent common ancestor of the Mediterranean *Tethysbaena* species—*T. ophelicola* from the coastal plain of Israel, *T. scabra* from the Balearic Islands, *T. ledoyeri* from Southern France, and *T. argentarii* from Italy—dated to the Tortonian in the Mid Miocene, 10.71 MYA (with 95% HPD of 6.27 – 6.04) as was previously found by Cánovas et al. (2016). The Ayyalon cave *Tethysbaena*, *T. ophelicola*, separated from other Mediterranean species around that time, 9.46 MYA (with 95% HPD of 14.20–5.71). The thermal water of the Ayyalon cave complex is part of the Yarkon-Taninim aquifer (Weinberger, Rosenthal, Ben-Zvi & Zeitoun, 1994). During Oligocene-Miocene regressions, canyons were entrenched along the Mediterranean Sea shoreline, serving as major outlets of the Yarkon-Taninim aquifer, potentially forming anchialine karst caves (Frumkin, Dimentman & Naaman, 2022; Laskow, Gendler, Goldberg, Gvirtzman & Frumkin, 2011). Page et al. (2016) hypothesized that the ancestral habitats of Thermosbaenacea are Tethyan anchialine caves. Accordingly, we can assume that the ancestor of *T. ophelicola* inhabited coastal anchialine caves in the Miocenic Tethys.

The most recent common ancestor of the Mediterranean and the Arabian-Caribbean clades of *Tethysbaena* is dated to the upper Eocene. During that period, the collision between the Arabian Plate and the Eurasian Plate resulted in the uplift of the Zagros Mountains in Iran (Mouthereau, Lacombe & Vergés, 2012). These mountain ranges acted as barriers, further isolating the Arabian Sea from the Mediterranean region (Sanmartín, 2003). The oldest, root node (*Tethysbaena*-*Halosbaena*) dated to 79.96 MYA (with the caveat of a low posterior probability and a large 95% HPD interval, 137.8 – 32.68 MYA). Page et al. (2016) established the phylogeny and divergence dates of the thermosbaeancean *Halosbaena*. They used the *Tethysbaena*-*Halosbaena* divergence as a calibration node, based on the presence of a continuous band of ocean crust through the length of the North Atlantic, indicating the maximum extent of the Tethys and the final opening of the Atlantic, dated to 107.5 MYA (with 95% HPD of 125–90). Thus, *Tethysbaena* ancestor in both our analysis and in Page et al. (2016) dates to the Cretaceous. The validity of the Paleozoic Ophel-driven hypothesis is also undermined by the deep phylogeny of peracaridean orders based on the small-subunit (SSU) rRNA gene, which showed that the thermosbaenacean lineage does not occupy a basal position relative to other peracarids (Spears, DeBry, Abele & Chodyla, 2005).

Overall, the molecular clock-based divergence patterns presented here do not support the previously proposed hypotheses regarding the origins of the Levantine *Tethysbaena*. Instead, we infer a complex, two-stage colonization pattern of the *Tethysbaena* species in the Levant: (1) a late Oligocene transgression event, through a marine gulf extending from the Arabian Sea in the East to the Sea of Galilea in the west, leading to the colonization of *T. relicta* in the Dead Sea-Jordan Rift Valley, and (2) a Miocene transgression event in the Mediterranean region, carrying *T. ophelicola* to the coastal plain of Israel. Our results also show that the Cretaceous *Tethysbaena* ancestor first established in present-day Morocco, and then diverged into two groups. The first is a Tethyan group including Oman, the Dead Sea-Jordan Rift Valley and the Caribbean Sea. The second group formed around the emerging Mediterranean Sea, in its marginal aquifers, including Ayyalon, Southern France, Italy and the Balearic Islands.

## CONCLUSIONS

Our results reject the morphology-based cladistics and suggest that *T. relicta* shared a most recent common ancestor with *Tethysbaena* species from Oman and Dominican Republic, whereas the circum-Mediterranean species, including *T. ophelicola*, shared another ancestor. The molecular dating analysis suggest a two-stage colonization of the *Tethysbaena* species in the Levant, explaining their distant origins: a late Oligocene transgression leading to the colonization of *T. relicta* in the Dead Sea-Jordan Rift Valley, and a Miocene transgression in the Mediterranean region followed by a marine regression, stranding *T. ophelicola* in the coastal plain of Israel. The speciose *Tethysbaena* provides an exquisite opportunity for testing paleogeographic paradigms. Here we analyzed the phylogenetic relationships and divergence of nine out of twenty-seven known *Tethysbaena* species using the mitochondrial barcode gene. Future studies should examine additional species utilizing more genes or complete genomes to further unveil the phylogeny and biogeography of this unique group of ancient subterranean crustaceans.

The study of these subterranean species is not only an opportunity to broaden our understanding of paleogeography; it is also paramount for the protection of the hidden biodiversity found in these largely inaccessible habitats, but which is nonetheless hugely influenced by human activity. Extraction of groundwater for irrigation and other uses, pollution, as well as quarrying, mining, and above-ground development may put these underground ecosystems at severe risk. The unique and often endemic nature of stygobiont species makes them even more prone to extinction, and extensive exploration of this under-explored biome, worldwide, is necessary in order to gain understanding and appreciation of the hidden biodiversity underground – an understanding that may pave the way for conservation of these species and their ecosystems.

## ACKNOWLEDGEMENTS

We are immensely grateful to Boaz Langford, Israel Naaman, Yoav Negev, Lior Enmar, Ilia Kutuzov, Shlomit Cooper-Frumkin, Amitai Cooper, and the Israel Cave Research Center team for their support in field sampling. We thank Chanan Dimentman for sharing invaluable knowledge on subterranean Levantine fauna, and Stas Malavin for providing helpful comments on the draft.

## DATA AVAILABILITY STATEMENT

The data underlying this article are available in the GenBank Nucleotide Database at https://www.ncbi.nlm.nih.gov/genbank/, and can be accessed with accession numbers OR189199–OR189204.

